# Endogenous esterases of *Clostridium thermocellum* are identified and disrupted for enhanced isobutyl acetate production from cellulose

**DOI:** 10.1101/761833

**Authors:** Hyeongmin Seo, Preston N. Nicely, Cong T. Trinh

## Abstract

Medium chain esters are potential drop-in biofuels and versatile chemicals. Currently, these esters are largely produced by the conventional chemical process that uses harsh operating conditions and requires high energy input. Alternatively, the microbial conversion route has recently emerged as a promising platform for sustainable and renewable ester production. The ester biosynthesis pathways can utilize either esterases/lipases or alcohol acyltransferase (AAT), but the AAT-dependent pathway is more thermodynamically favorable in aqueous fermentation environment. Even though cellulolytic thermophiles such as *Clostridium thermocellum* harboring the engineered AAT-dependent pathway can directly convert lignocellulosic biomass into esters, the production is currently not efficient and requires optimization. One potential bottleneck is the ester degradation caused by the endogenous carbohydrate esterases (CEs) whose functional roles are poorly understood. In this study, we developed a simple, high-throughput colorimetric assay to screen the endogenous esterases of *C. thermocellum* responsible for ester hydrolysis. We identified, characterized, and disrupted two critical endogenous esterases that significantly contributes to isobutyl acetate degradation in *C. thermocellum*. We demonstrated that not only did the engineered esterase-deficient strain alleviate ester hydrolysis but also helped improve isobutyl acetate production while not affecting its robust metabolism for effective cellulose assimilation.

**IMPORTANCE:** Carbohydrate esterases (CEs) are important enzymes in the deconstruction of lignocellulosic biomass by the cellulolytic thermophile *C. thermocellum*, yet some are potential ester degraders in a microbial ester production system. Currently, the functional roles of CEs for hydrolyzing medium chain esters and negatively affecting the ester microbial biosynthesis are not well understood. This study discovered novel CEs responsible for isobutyl acetate degradation in *C. thermocellum* and hence identified one of the critical bottlenecks for direct conversion of lignocellulosic biomass into esters.

## INTRODUCTION

Medium chain (C_6_-C_10_) esters are potential drop-in biofuels and valuable chemicals with broad industrial applications such as food additives, fragrances, and solvents (1-4). Isobutyl acetate, for instance, is a biodegradable solvent that resembles methyl isobutyl ketone and toluene in many formulations. Esters are commonly produced by the conventional Fisher esterification process that condenses a carboxylic acid and an alcohol in aqueous environments with requirement of heat and strong acid. This process, however, is not environmentally friendly and thermodynamically favorable where additional energy input is required to remove water, a byproduct of the esterification, to avoid inhibition during the reaction (5).

The microbial ester production has recently emerged as an alternative promising platform for ecofriendly biosynthesis of esters from renewable and sustainable feedstocks (6-9). The ester biosynthesis pathways can utilize either esterases/lipases (10) or alcohol acetyltransferases (AATs) (7, 8). Unlike the esterase/lipase-dependent pathway that follows the Fisher esterification chemistry, the AAT-dependent pathway makes esters by condensing an alcohol and an acyl-CoA. This reaction is thermodynamically favorable in aqueous fermentation conditions because water is not generated as a byproduct and the stored energy from the thioester bond of acyl-CoA helps effective transfer of an acyl group. This thermodynamic advantage of the AAT-dependent pathway makes it attractive for ester biosynthesis in microorganisms (6-9, 11).

Among microbes, cellulolytic bacteria have increasingly become attractive for direct conversion of lignocellulosic biomass into esters in a consolidated bioprocessing (CBP) where biomass degradation and fermentation simultaneously take place in a single step and hence reduce the production cost (12). The gram-positive thermophilic anaerobe *Clostridium thermocellum* is an ideal cellulolytic bacteria for CBP because it is the best biomass degrader known to date and has endogenous metabolic pathways for synthesizing various alcohols including ethanol and isobutanol (13-15). While native *C. thermocellum* cannot produce esters, we have recently demonstrated by expressing a thermostable chloramphenicol acetyltransferase (CAT) in *C. thermocellum*, the direct biosynthesis of medium chain esters from cellulose at elevated temperature (55°C) is feasible (16).

The current ester production in *C. thermocellum* is not efficient and requires optimization. One of the potential bottlenecks is the ester degradation caused by endogenous esterases. For example, by disrupting an esterase that hydrolyzes volatile esters from *Saccharomyces cerevisiae* (17, 18), biosynthesis of medium chain esters was increased (19), suggesting that balance of AAT and esterase activity of brewer yeasts is important for the ester accumulation in wine and distillates (19-21). Like many cellulolytic bacteria, *C. thermocellum* possesses multiple carbohydrate esterases (CEs) whose expression helps deconstruct lignocellulosic biomass in a concerted enzyme reaction (22-24). Currently, it is widely unknown about the functional roles of CEs in ester degradation in *C. thermocellum*.

In this study, we identified, characterized, and disrupted endogenous esterases responsible for the isobutyl acetate degradation in *C. thermocellum*. We started by developing a high-throughput screening platform to screen for esterases responsible for ester hydrolysis. Next, we identified and disrupted the critical esterases that helped alleviate the isobutyl acetate degradation in *C. thermocellum* while not affecting cellulose utilization. Finally, we demonstrated the engineered esterase-deficient *C. thermocellum* strain improved isobutyl acetate production.

## RESULTS AND DISCUSSION

### Thermodynamic analysis suggests favorable hydrolysis of isobutyl acetate by an esterase under aqueous fermentation environment

The driving force for the esterase-dependent pathway to synthesize esters depends on the equilibrium constant and species concentrations (Figure 1A). For isobutyl acetate biosynthesis in an aqueous solution, the equilibrium constant (K_eq_ =1.3×10^−5^ M^-1^) at standard conditions (T = 37°C, P = 1 atm, pH 7, [C] = 1M) is relatively low, yielding a positive standard ΔG° of 27.8 ± 4.2 kJ/mol (25). This reaction is not thermodynamically favorable for ester biosynthesis but degradation (or hydrolysis in the reverse direction) and therefore, relatively high substrate concentrations are required to push the reaction forward (Table S1). In contrast, the AAT-dependent pathway thermodynamically favors the forward reaction (Figure 1B) with K_eq_ of 2.42 M^-1^ and a negative standard ΔG° of −2.4 ± 4.1, kJ/mol) (25), hence only relatively low substrate concentrations are needed for ester biosynthesis to occur (Table S1). Indeed, *C. thermocellum* expressing a heterologous AAT has recently been demonstrated to produce isobutyl acetate from lignocellulosic biomass (Figure 1C) (16). Overall, esterases play a critical role in ester degradation under aqueous fermentation conditions while AATs are essential for ester biosynthesis.

**Figure 1.**
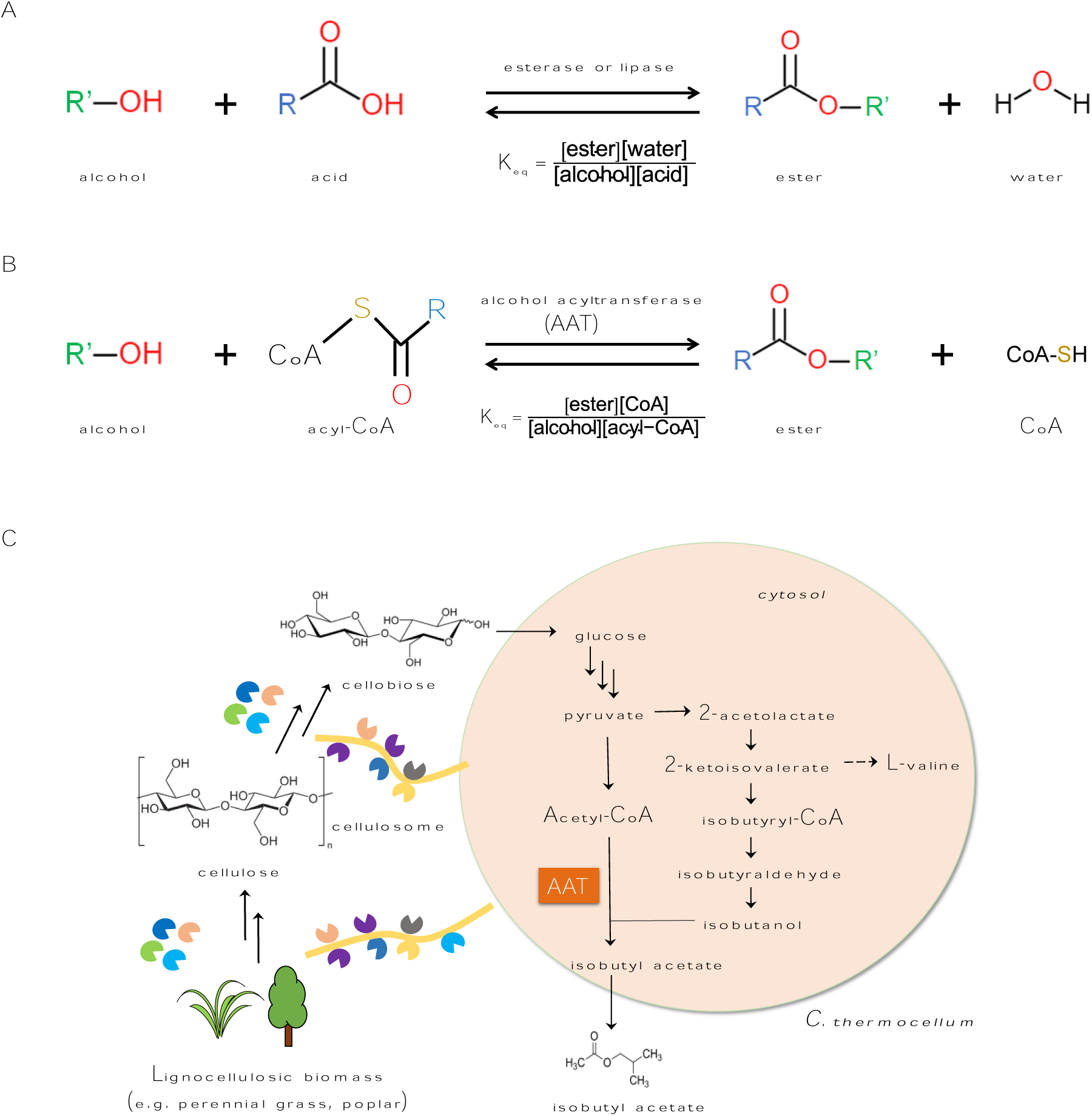
Metabolic pathways for isobutyl acetate biosynthesis in *C. thermocellum*. **(A)** The esterase/lipase-dependent ester biosynthesis pathway. **(B)** The alcohol acyltransferase (AAT)-dependent ester biosynthesis pathway. **(C)** *De novo* isobutyl acetate biosynthesis directly from lignocellulosic biomass via the AAT-dependent pathway in *C. thermocellum.*

### A developed high-throughput screening enabled identification of two esterases hydrolyzing isobutyl acetate

Even though the AAT-dependent pathway is shown to be feasible in *C. thermocellum* (16), the production is inefficient. One of the potential bottlenecks might be due to relatively high endogenous activity of esterases in *C. thermocellum* that limit the ester biosynthesis. Even though *C. thermocellum* harnesses a large number of CEs to deacetylate O-acetyl groups of hemicelluloses responsible for lignocellulosic biomass deconstruction (22-24), their functional roles for ester degradation are largely unknown. Therefore, it is critical to identify CEs hydrolyzing isobutyl acetate in *C. thermocellum* and understand their effects on the isobutyl acetate biosynthesis from lignocellulosic biomass during fermentation.

We started by searching CEs of *C. thermocellum* in the CAZy database (26). We found that *C. thermocellum* has 14 CEs that are classified as either N-deacetylase or O-deacetylase, based on the domains and putative substrates of CEs (Table S2). Since isobutyl acetate is an O-acetyl ester that can be deacylated by O-acetyl esterases, we focused on investigating the seven putative O-acetyl esterases (Table 1). Based on the sequence motifs, only Clo1313_0613 is a cytosolic enzyme while others are likely secreted and/or anchored to the cell membrane (Figure 2A). To characterize these seven CEs for ester hydrolysis, we heterologously expressed them in *E. coli*. We first removed the signal peptides of these proteins except Clo1313_0613 and tagged all proteins with 6X His residues at their N-termini (Figure 2A, Table 2). In the case of Clo1313_1424, we truncated it to contain one Xyn_B like domain instead of the duplicated domain present in the original sequence. Successful expression of the seven modified esterases in *E. coli* was confirmed (Figure 2B).

**Table 1.**
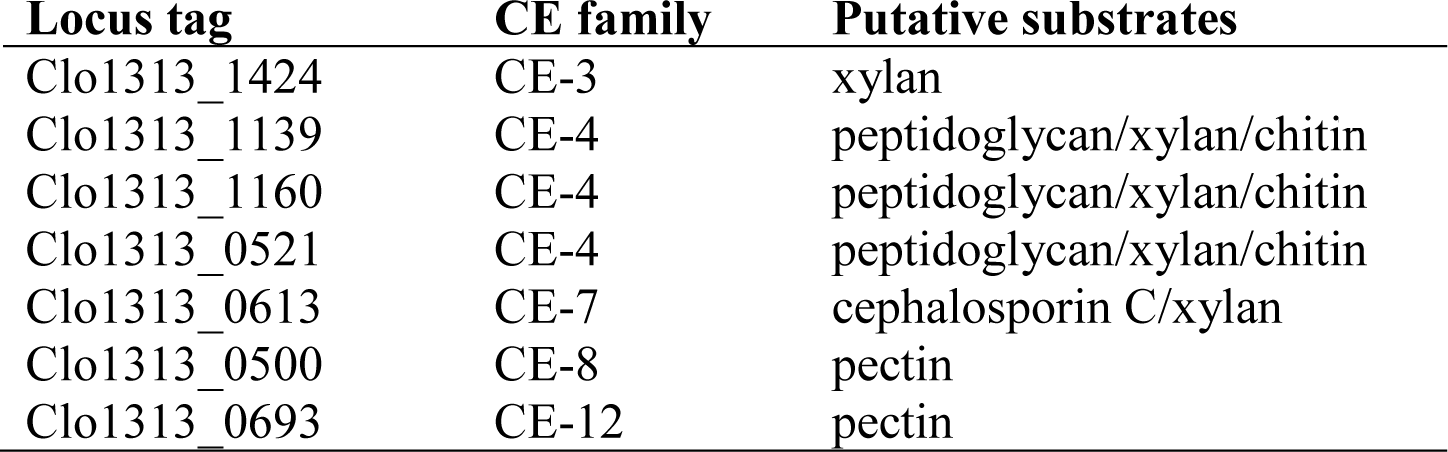
Seven O-deacetylases and their putative substrates.

**Table 2.**
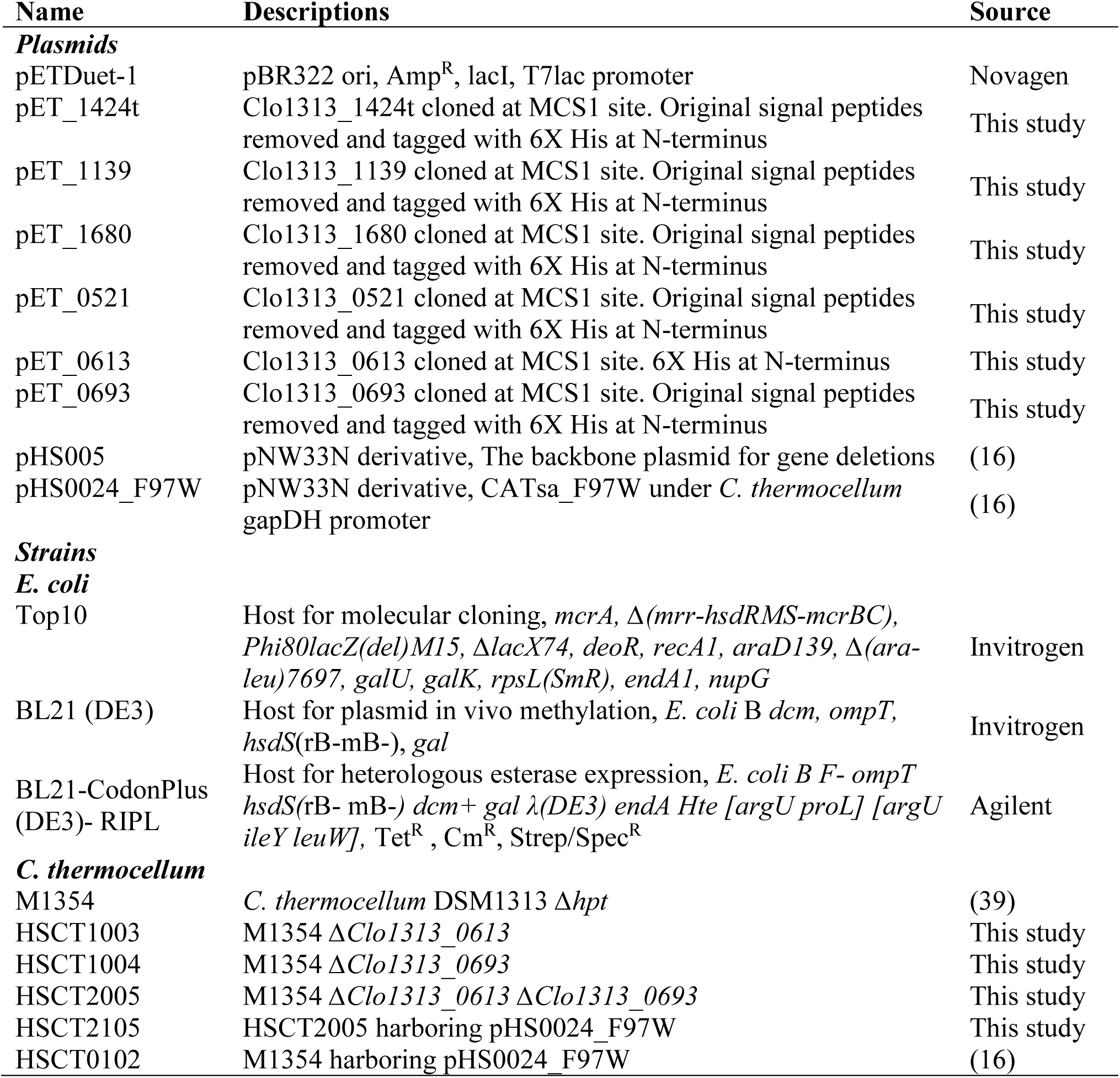
Plasmids and strains used in this study.

**Figure 2.**
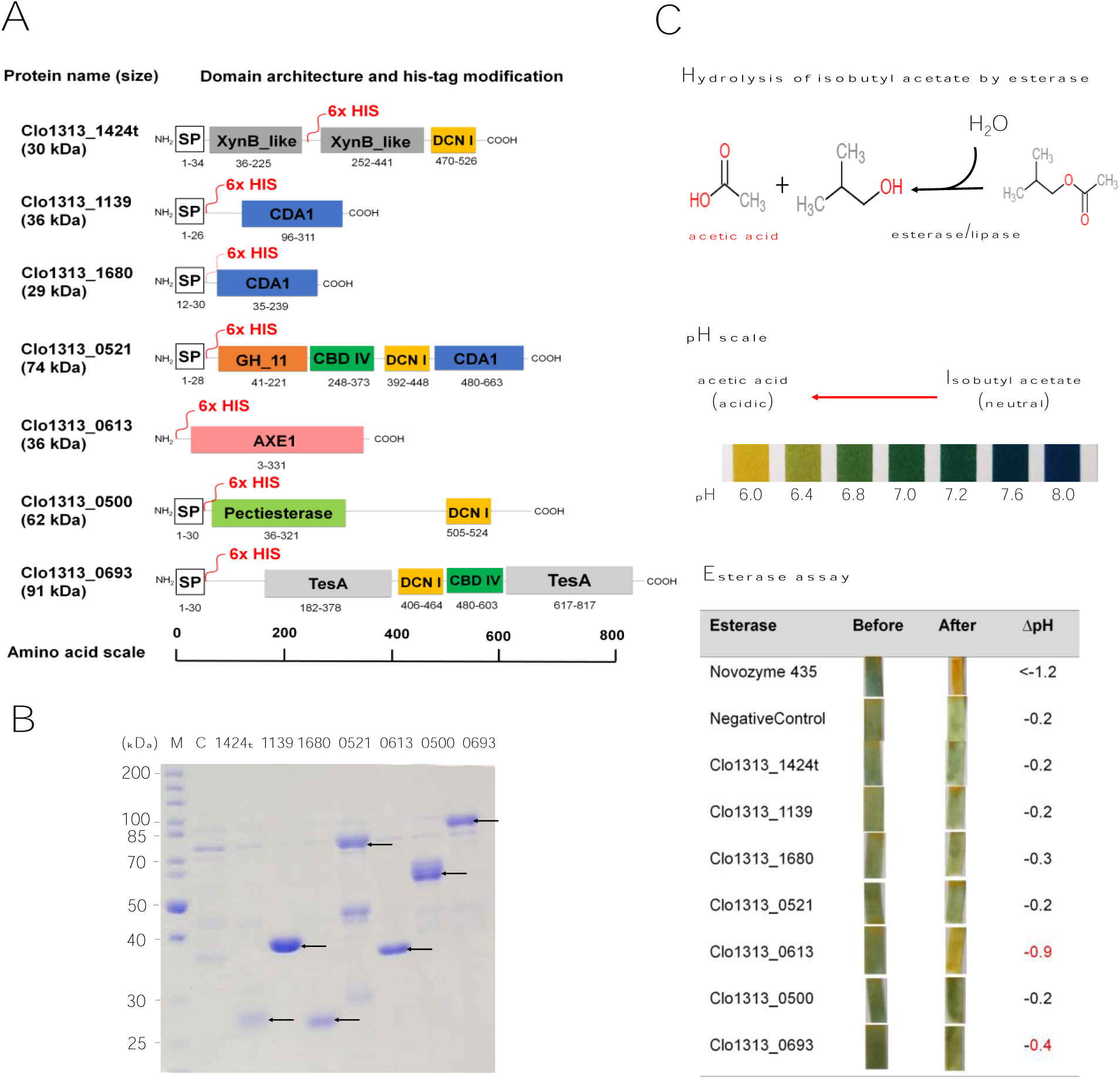
Identification of esterases responsible for isobutyl acetate hydrolysis in *C. thermocellum*. **(A)** Seven O-deacetylases in *C. thermocellum* and their domain architecture. The protein annotations followed their gene locus tag. The locations tagged with the histidine residues were marked as 6x HIS in red. Clo1313_1424t is a truncated version of Clo1313_1424 which is derived from removal of the signal peptide and one XynB_like domain located at 36-225 residues that is identical to the other located at 252-441 residues. Abbreviation: SP; signal peptide, DCN I; dockerin I. **(B)** SDS-PAGE of his-tag purified proteins expressed in the recombinant *E. coli* strains. Abbreviations: M: protein marker ladder; C: *E. coli* harboring an empty backbone plasmid. **(C)** A high-throughput, colorimetric esterase assay using the pH strips. The ΔpH values represent an average pH change of three biological replicates.

To evaluate their activities toward isobutyl acetate hydrolysis, we developed a simple, high throughput method based on pH change (Figure 2C). The hydrolysis of isobutyl acetate (pKa ∼25) by an esterase into isobutanol and acetic acid (pKa ∼4.75) lowers pH that can be determined by a colorimetric assay. We screened the esterase activity towards isobutyl acetate by pH change upon enzymatic reactions with crude extracts of the esterase-overexpressing *E. coli* strains. As a positive control, the reaction with Novozyme 435, an immobilized CalB lipase, resulted in a dramatic pH change, validating that the hydrolysis of isobutyl acetate lowers pH. We found that only Clo1313_0613 and Clo1313_0693 out of the seven CEs characterized could hydrolyze isobutyl acetate with pH changes of −0.9 and −0.4, respectively. The hydrolysis was further confirmed with detection of isobutanol and acetate by HPLC analysis. For the negative control, we observed that the reaction without expressed heterologous esterase slightly lowered pH (ΔpH = −0.2), likely due to either endogenous esterase activity of *E. coli* (8) or spontaneous hydrolysis in the aqueous environment. However, the hydrolysis in the negative control experiment was very low as detection of acetate and isobutanol by HPLC analysis was unquantifiable. Taken together, we identified the two esterases, Clo1313_0613 and Clo1313_0693, that can hydrolyze isobutyl acetate.

### Homolog proteins of the identified esterases widely exist in bacteria regardless of their cellulolytic capability

We next questioned whether the two identified esterases are only present in the cellulolytic bacteria such as *C. thermocellum* for their contributions to biomass deconstruction. Bioinformatic analysis revealed that Clo1313_0613 (cephalosporin C deacetylase) shares 59.09% protein sequence identity with a well-studied homo-hexamer cephalosporin C deacetylase from *Bacillus subtilis* (Figure S1). This deacetylase has a broad substrate range with high specificity towards short substrates including p-nitrophenyl acetate (pNPA) and short chain or pre-processed xylans (27). Our *in vitro* assay indeed confirmed that Clo1313_0613 exhibited activity towards pNPA (Figure S2). Based on the constructed phylogenetic tree of Clo1313_0613, we found homolog proteins exist in a wide spectrum of bacteria, regardless of their cellulolytic capability (Figure 3A). Most of the bacteria identified were gram-positive including *Actinomyces* sp. while there was a small group of gram-negative bacteria also found such as *Mesorhizobium sp*. Detailed sequence analysis shows that Clo1313_0613 and homolog proteins possess highly conserved active sites with GxSxG motif (Figure S3A), which is present in most esterases and lipases (28). Taken together, Clo1313_0613 is an intracellular esterase with a broad substrate range and has little relationship with cellulose metabolism. Due to the broad substrate range, the biological role of this enzyme remains unclear and its substrate specificity needs further examination for deeper understanding.

**Figure 3.**
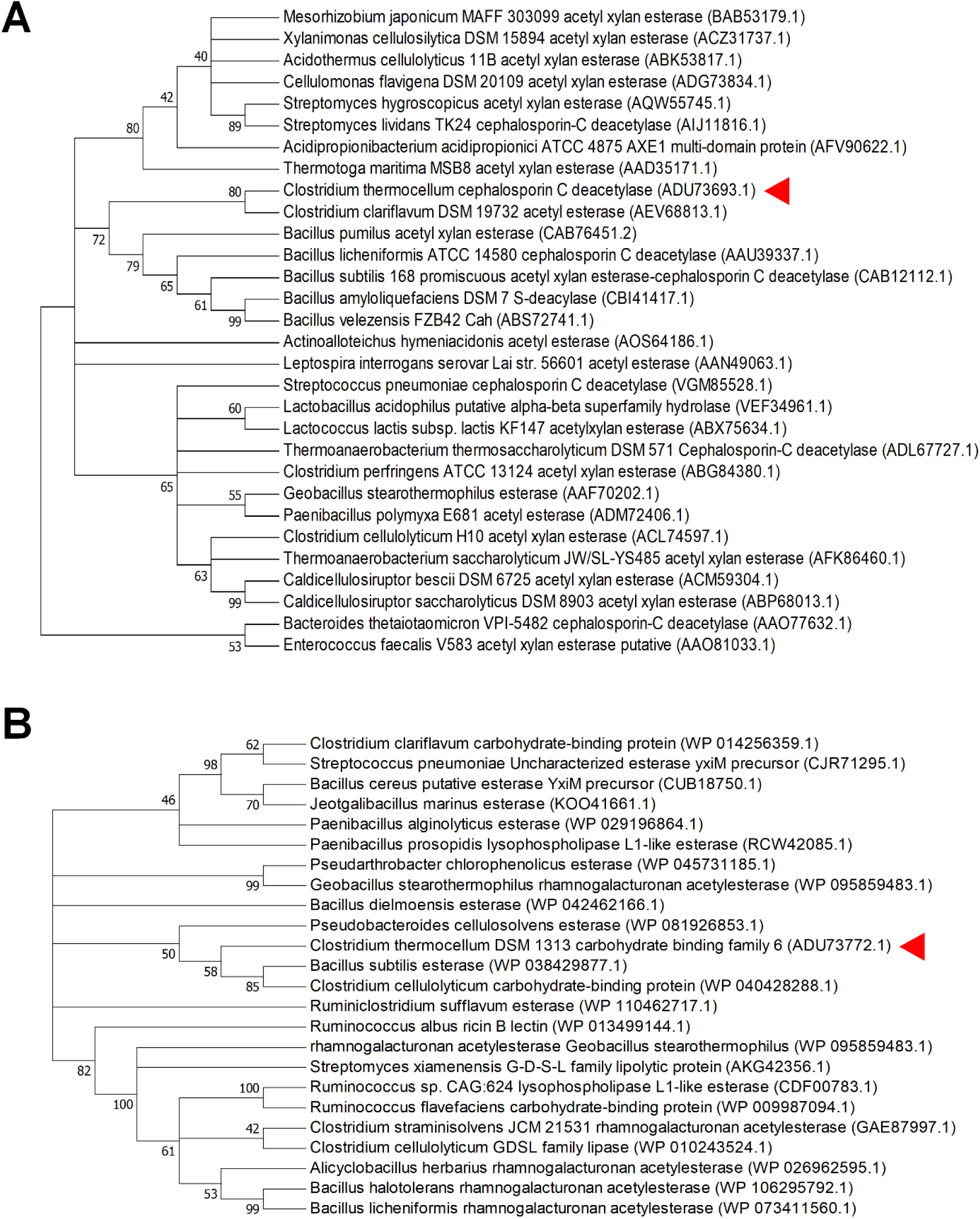
Phylogenetic analysis of Clo1313_0613 and Clo1313_0693. **(A)** A phylogenetic tree of AXE1 domain in Clo1313_0613. The red triangle indicates Clo1313_0613. **(B)** A phylogenetic tree of TesA domain in Clo1313_0693. The red triangle indicates Clo1313_0693.

Clo1313_0693, belonging to the carbohydrate binding family 6, shares the highest sequence similarity (67.65%) with an uncharacterized protein YxiM from *B. subtilis* (locus tag: BSU_39120). The YxiM protein has not been experimentally characterized, but its homolog proteins deacetylate broad substrates including apple pectin rhamnogalacturonan (29), pNPA (29, 30) and cephalosporin C (30). Clo1313_0693 possesses two rhamnogalacturan acetylesterase-like domains that are likely responsible for the hydrolysis activity of isobutyl acetate as experimentally observed. From the constructed phylogenetic tree of Clo1313_0693, we found that homolog proteins broadly exist in both cellulolytic and non-cellulolytic bacteria (Figure 3B), suggesting that Clo1313_0693 might not be critical for cellulose metabolism. Clo1313_0693 and the homolog proteins exhibit catalytic triads (Ser-His-Asp) at active sites (Figure S3B), which is very different from Clo1313_0613. The nucleophile, serine, is located near N-terminus whereas histidine and aspartate are located near C-terminus of the domain. These features are highly correlated with rhamnogalacturonan acetylesterases (31) that are involved in pectin deconstruction. Especially, the gene encoding Clo1313_0693 is located under a promoter regulated by σ_I3_ RNA polymerase (32, 33), which is related to a membrane-associated anti-sigma factor (Rsigl3) that specifically binds to pectin (34). Another pectinesterase under the σ_I3_ regulon is Clo1313_1983, a rhamnogalacturonan lyase (33). Note that Clo1313_0693 includes a signal peptide and dockerin (Figure 2A), which is proposed to be a component of extracellular cellulosome (35).

In summary, the identified esterases hydrolyzing isobutyl acetate in *C. thermocellum* are evolutionarily distinct and have homologous proteins in multiple bacteria including non-cellulolytic microbes.

### Disruption of the esterases from *C. thermocellum* alleviates hydrolysis of isobutyl acetate

Transcriptomic analysis of *C. thermocellum* shows that both of the esterases identified are expressed in *C. thermocellum* (35, 36). To test their *in vivo* activities for isobutyl acetate degradation, we disrupted these esterase genes through homologous recombination (Figure S4A). We generated three esterase-deficient strains including HSCT1003 with Clo1313_0613 deletion, HSCT1004 with Clo1313_0693 deletion, and HSCT2005 with both Clo1313_0613 and Clo1313_0693 deletion (Figure 4A, Table 2). Growth characterization shows that HSCT1003 (0.271 ± 0.017 1/h) and HSCT2005 (0.270 ± 0.031 1/h) grew about 20% slower than the parent strain (M1354, 0.341 ± 0.013 1/h) while HSCT1004 (0.352 ± 0.018 1/h) did not exhibit significant change in growth rate. Interestingly, the cellulose degradation profiles were similar among the engineered strains (Figure 4B). Therefore, we concluded that the esterase disruptions did not inhibit cellulose consumption in *C. thermocellum*.

**Figure 4.**
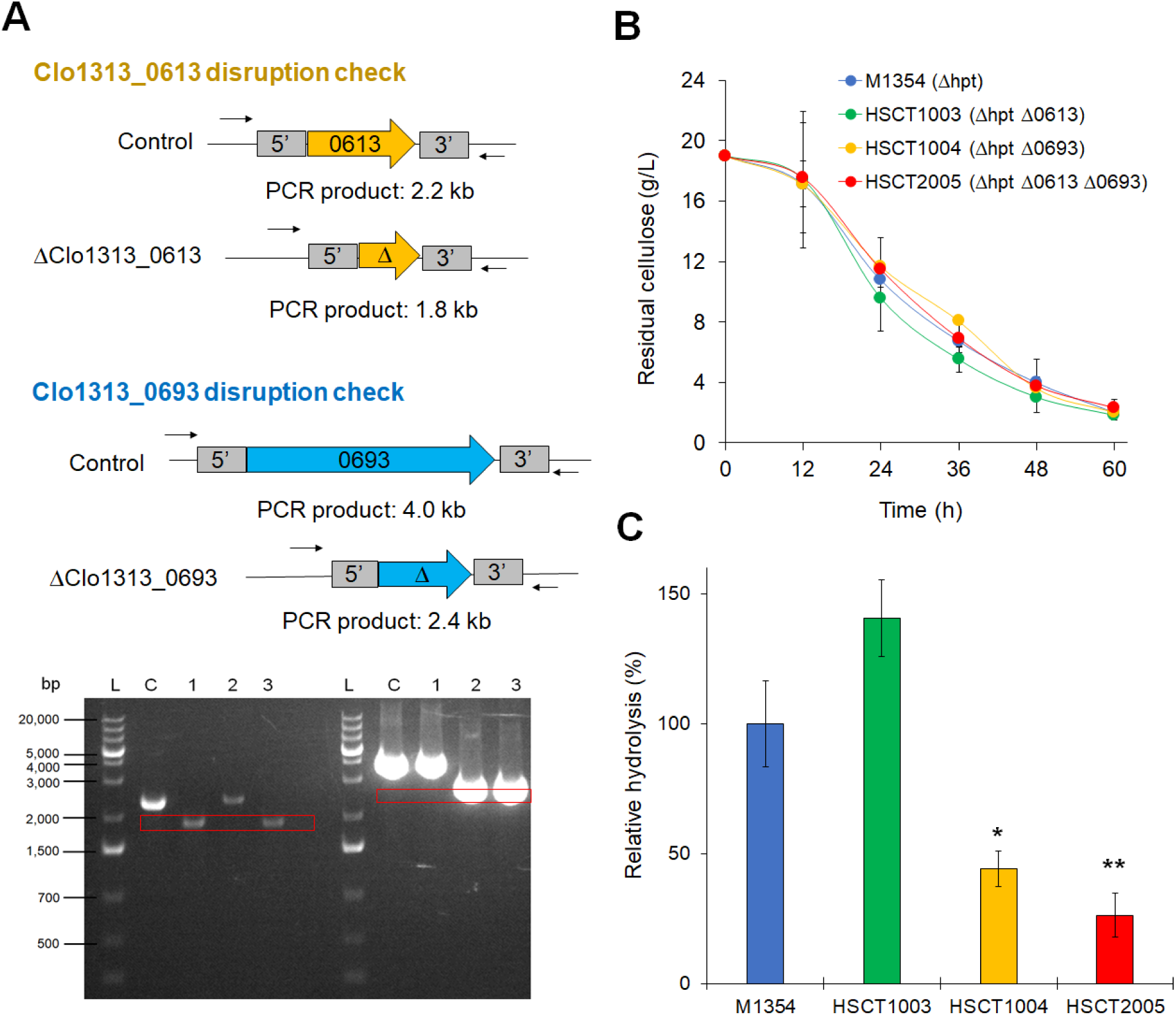
Disruption of esterases in *C. thermocellum* and characteristics of the mutants. **(A)** Confirmation of gene disruption. Abbreviations: L: DNA marker ladder; C: control strain (M1354); 1: HSCT1003 (ΔClo1313_0613); 2: HSCT1004 (ΔClo1313_0693); 3: HSCT2005 (ΔClo1313_0613, ΔClo1313_0693). **(B)** Comparison of residual cellulose by the engineered *C. thermocellum* strains during 20 g/L cellulose fermentation. The error bar represents standard deviation of three biological replicates. **(C)** Relative hydrolysis of isobutyl acetate by the engineered strains. Statistical analysis: t-test; “*”: p-value < 2.3×10^−4^ (t = 6.937, df = 7); “**”: p-value < 4.8×10^−5^ (t= 8.862, df = 7). The error bars represent standard deviation of six biological replicates.

We next compared the hydrolysis of isobutyl acetate among the parent and esterase-deficient strains (Figure 4C). To estimate the relative *in vivo* esterase activity, we externally supplemented 1 g/L (8.61 mM) of isobutyl acetate and measured isobutanol released by HPLC. HSCT1003 showed no change in isobutyl acetate hydrolysis while HSCT1004 reduced isobutyl acetate degradation by 56% as compared to the parent strain (Figure 4C). Interestingly, a combination of esterase deletion in HSCT2005 significantly reduced isobutyl acetate hydrolysis (0.57 mM) by 75%.

Taken together, the identified two esterases were successfully disrupted, and the genetic modification in *C. thermocellum* alleviated hydrolysis of isobutyl acetate. This result is consistent with the *in vitro* activity of esterases observed.

### Engineered esterase-deficient *C. thermocellum* improved isobutyl ester production

In our previous study, we developed a thermostable ester production module, harboring an engineered thermostable CAT_Sa_ F97W under a strong gapDH promoter, that is capable of producing esters directly from cellulose in *C. thermocellum* at elevated temperature (T ≥ 50°C) (16). To evaluate whether the alleviated esterase activity improved isobutyl acetate production in *C. thermocellum*, we introduced the thermostable ester production module into HSCT2005 to generate the esterase-deficient, ester-producing strain HSCT2105. Strain characterization showed that HSCT2105 produced 3.1 mg/L of isobutyl acetate (Figure 5A) after the 72-hour cellulose fermentation. In comparison to HSCT0102 (Figure 5B), harboring the identical production module without the esterase disruption, HSCT2105 achieved 1.6-fold higher titer (16). Since both strains yielded the same maximal pellet protein (∼520 mg/L) and consumed cellulose (∼17 g/L), it further supports that the esterase disruptions did not interfere with cellulose utilization.

**Figure 5.**
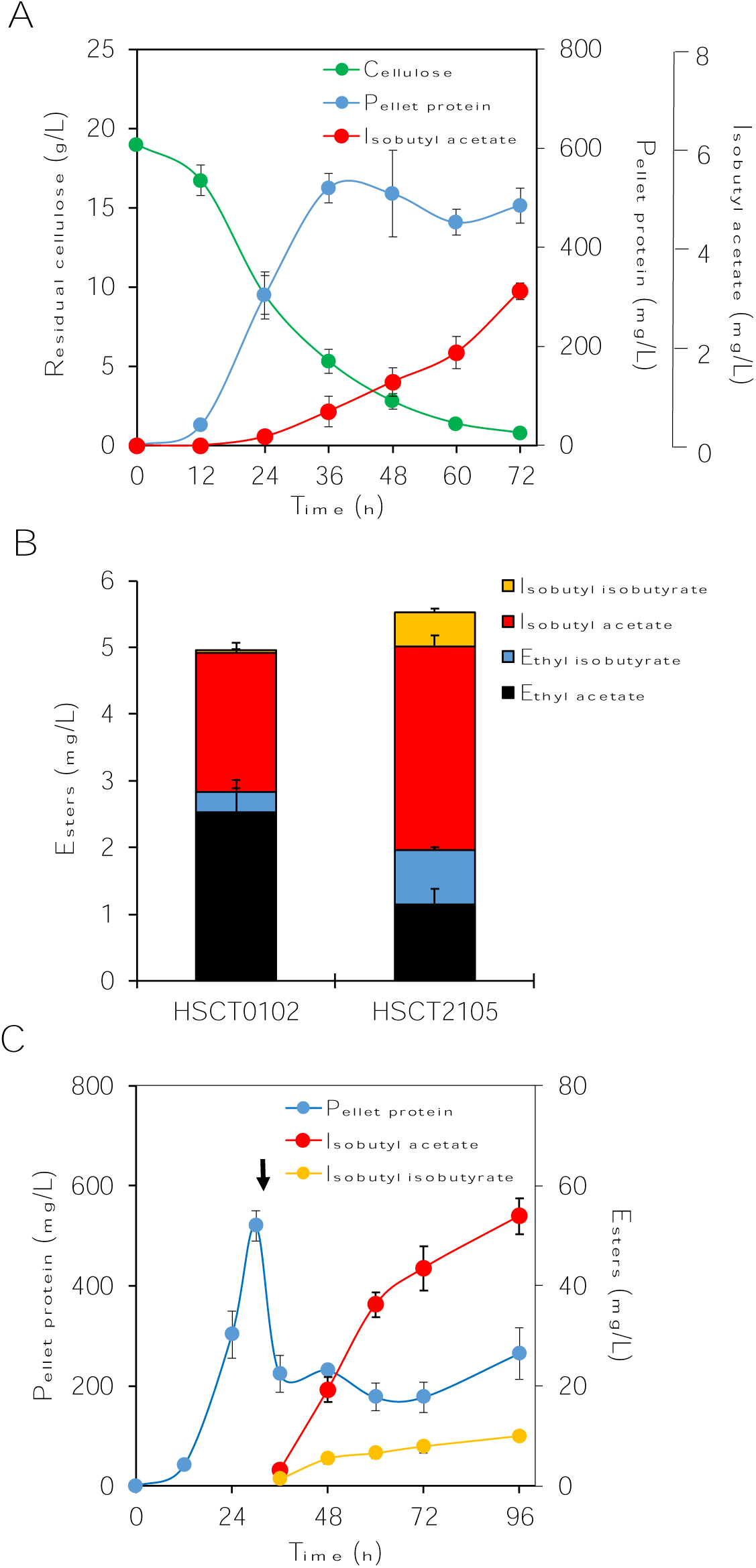
Ester production by HSCT2105. **(A)** A kinetic profile of isobutyl acetate production from cellulose by HSCT2105. **(B)** Comparison of ester production by HSCT0102 and HSCT2105. Esters were quantified after 72-hour cellulose fermentation. **(C)** A kinetic profile of pellet protein and isobutyl ester concentration in the cellulose fermentation with external supply isobutanol. The black arrow indicates the time point when 5 g/L isobutanol was added. The error bars indicate standard deviation of three biological replicates.

In addition, we observed that HSCT2105 produced two times more isobutyl-CoA related esters (i.e., ethyl isobutyrate, isobutyl isobutyrate) than HSCT0102 (Figure 5B). Since *C. thermocellum* has five native pyruvate ferredoxin oxidoreductase (PFOR) genes or gene clusters, these enzymes are likely responsible for converting 2-ketoisovalerate to isobutyl-CoA (14, 37). The increase in production of isobutyrate esters (i.e., isobutyl acetate, isobutyl isobutyrate and ethyl isobutyrate) suggest that HSCT2105 might have a higher metabolic flux to the isobutyl-CoA pathway than HSCT0102 has. This result correlates with a decrease in lactate production but an increase in isobutanol production (Table 3). Currently, we do not have a clear explanation of why the esterase disruptions increased the isobutyl-CoA flux.

**Table 3.**
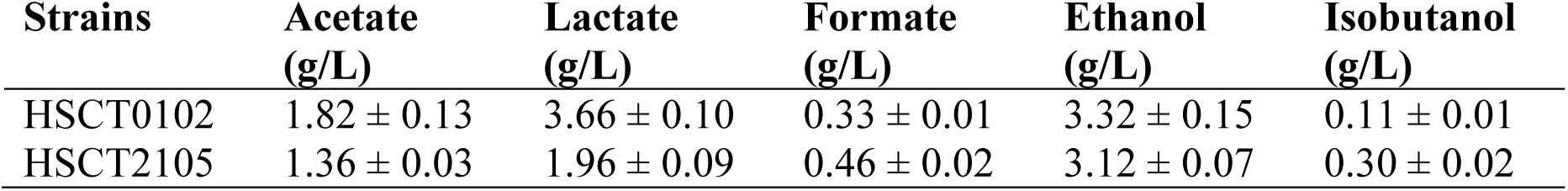
Endpoint fermentative metabolites of cellulose fermentation by HSCT0102 and HSCT2105. The values represent mean ± standard deviation from three biological replicates.

The observed increase in isobutyl-CoA flux motivated us to test whether isobutyl ester production (i.e., isobutyl acetate and isobutyl isobutyrate) can be further improved by supplementing external isobutanol (Figure 5C). Upon external supply of 5g/L isobutanol, pellet protein of HSCT2105 sharply dropped down to 220 mg/L, which might attribute to toxicity of isobutanol (38). As expected, we observed the increase in isobutyl acetate and isobutyl isobutyrate production of HSCT2105 up to 53.9 mg/L and 9.9 mg/L, respectively, after 66-hour cellulose fermentation.

Overall, the esterase-deficient, ester-producing *C. thermocellum* HSCT2105 showed a higher metabolic capacity for ester biosynthesis. It enhanced production of not only isobutyl acetate but also ethyl isobutyrate and isobutyl isobutyrate. This strain can serve as a starting host for further metabolic engineering to enhance ester production.

## CONCLUSION

Esterases are not only important for lignocellulosic biomass deconstruction but also detrimental for ester biosynthesis. Hence, the *in vivo* control of esterases presents one of the critical metabolic engineering targets for enhanced microbial ester production. Through bioinformatic analysis together with enzyme and strain characterization, we discovered critical esterases responsible for hydrolyzing isobutyl acetate in *C. thermocellum* using a simple, high-throughput, colorimetric screening assay developed. By deleting these esterase genes, we created an esterase-deficient, ester-producing *C. thermocellum* that improves ester biosynthesis while not affecting the strain capability for cellulose utilization. We anticipate the esterase-deficient *C. thermocellum* would serve as a useful thermophilic CBP platform for ester production.

## MATERIALS AND METHODS

### Strains and plasmids

Bacterial strains and plasmids used in this study are listed in Table 2. *C. thermocellum* DSM 1313 Δhpt (M1354) (39) was used as the parent strain because the hpt deletion allows counter-selection for markerless gene deletion without effecting cell growth (40). *Escherichia coli* TOP10 and BL21(DE3) were used for general molecular cloning and dam methylation of plasmids for *C. thermocellum* transformation (41), respectively. For heterologous expression of esterases, *E. coli* BL21 CodonPlus (DE3)-RIPL (Agilent, CA, USA) was used to avoid potential expression problems caused by codon bias.

### Chemicals and reagents

Unless noted elsewhere, all chemicals were purchased from Thermo Fisher Scientific (MA, USA) and/or Sigma-Aldrich (MO, USA). Restriction enzymes and T4 ligase were purchased from New England Biolabs (MA, USA).

### Media and cultivation

*E. coli* strains were grown in lysogeny broth (LB) medium supplemented with 30 µg/mL chloramphenicol (Cm) and/or 100 µg/mL ampicillin (Amp) when appropriate. All *C. thermocellum* strains were cultivated in an anaerobic chamber (Sheldon manufacturing, OR, USA) with an anaerobic gas mixture (90% N_2_, 5% CO_2_, 5% H_2_) or rubber stopper sealed Balch tubes outside the chamber. For genetic engineering, *C. thermocellum* strains were grown in rich media, CTFuD or CTFuD-NY (40) supplemented with 10 µg/mL thiamphenicol (Tm) as needed. The CTFuD medium contained 2 g/L yeast extract while CTFuD-NY contained vitamins and trace elements (40) without the yeast extract. Except for the isobutyl acetate production from cellulose, *C. thermocellum* strains were grown in a defined MTC medium with 5 g/L cellobiose as previously described (42). To produce isobutyl acetate from cellulose, a modified MTC medium (C-MTC) was used as previously described (16). The C-MTC medium contained (per liter): 2 g urea, 1.5 g ammonium chloride, 10 g 3-(N-morpholino)propanesulfonic acid (MOPS), 2 g sodium citrate tribasic dihydrate, 1.5 g citric acid monohydrate, 1 g sodium sulfate, 0.5 g potassium phosphate monobasic, 1 g cysteine-HCl, 0.2 g calcium chloride, 1 g magnesium chloride hexahydrate, 0.1 g iron (II) chloride tetrahydrate, 2.5 g sodium bicarbonate, 0.02 g pyridoxamine dihydrochloride, 0.004 g p-aminobenzoic acid, 0.002 g biotin, and 0.002 g vitamin B12. The medium was adjusted to pH of 7.5 and autoclaved for sterilization. Solid LB and CTFuD media additionally contained 15 g/L and 10 g/L of agar, respectively.

### Molecular cloning

#### Plasmid construction

The primers used in this study are listed in Table S3. Plasmids were constructed by ligation-dependent cloning and/or Gibson DNA assembly. Briefly, DNA fragments were amplified using the Phusion DNA polymerase (cat# F530S, Thermo Fisher Scientific, MA, USA) and then purified by DNA purification and gel extraction kits (Omega biotek, GA, USA). For the ligation-dependent cloning, the vectors and inserts were digested by restriction enzymes and ligated together using a T4 DNA ligase. In the case of Gibson DNA assembly cloning, the purified DNA fragments of the vector and insert were mixed together with the Gibson master mix (43) and assembled in 50°C for one hour. Using the DNA mixtures, *E. coli* TOP10 was transformed by heat-shock transformation and selected on LB agar plates (15 g/L agar) with appropriate antibiotics. All the constructed plasmids were checked by PCR amplification and/or restriction enzyme digestion, and Sanger sequencing.

#### C. thermocellum transformation

*C. thermocellum* cells were transformed by electroporation (16, 40). Electroporation was performed outside of the anaerobic chamber followed by cell recovery and plating inside the chamber. A series of two consecutive exponential pulses were applied using the electroporation system (cat # 45-0651, BTX Technologies Inc., MA, USA) set at 1.8 kV, 25 μF, and 350 Ω, which usually resulted in a pulse duration of 7.0-8.0 ms.

#### Markerless gene disruption

Markerless gene deletion was carried out by multiple selection steps (Figure S4A) (40). pHS005 (Figure S4B), derived from pNW33N, was used as a backbone plasmid (16). Each 500 to 750bp length upstream (5’) and downstream (3’) of the target gene were assembled together by the overlapping PCR method and cloned in the MCS1 of pHS005. In the MCS2, 500 to 750bp intermediate fragment (int) were cloned. The *dam* methylated plasmid was introduced to *C. thermocellum* and selected in CTFuD agar supplemented with Tm. A colony from successful transformation was selected and cultured in the liquid CTFuD-NY medium supplemented with Tm until the cell culture reached mid-log growth phase (OD ∼ 0.2-0.6). Cells were then serial diluted 100-fold and plated on a CTFuD-NY agar plate supplemented with Tm and 30 µg/mL 5-fluoro-2′-deoxyuridine (FuDR) followed by 3-5 days of incubation at 55°C. The colonies were PCR screened using the primers, HS252 and HS253 (Table S3), to check plasmid loss (Figure S4A). A colony without a PCR product was selected and streaked on the CTFuD-NY agar plate containing Tm and FuDR followed by incubation at 55°C for 3-5 days. Colonies were PCR screened again to ensure plasmid loss and then cultured in the liquid CTFuD-NY medium without selection. The cultured cells were serial diluted 100-fold and plated on the CTFuD-NY agar supplemented with 500 µg/mL 8-azahypoxanthine (8-AZH). After three days, colonies were PCR screened using the primers binding at genome-specific upstream and downstream sites of the target knockout gene (Table S3, Figure S4A). A colony was streaked on a new CTFuD-NY plate up to three transfers to isolate a single genotype. Finally, the markerless gene deletion was confirmed by negative cell growth in CTFuD medium supplemented with Tm.

### *In vitro* enzyme assays

#### Heterologous expression of esterases

To screen esterase activity towards isobutyl acetate, seven putative esterases from *C. thermocellum* were heterologously expressed in *E. coli* BL21 CodonPlus-RIPL(DE3) strains under a T7lac promoter. The recombinant *E. coli* strains were cultured overnight in 3 mL of LB medium at 37°C, transferred to 25 mL of fresh LB medium with 100-fold dilution, and cultured at 37°C until OD reached 0.3∼0.4. Then, protein expression was induced by adding isopropyl β-D-1-thiogalactopyranoside (IPTG) at final concentration of 0.1 mM. The IPTG-induced cells were cultured in a water bath orbital shaker (MaxQ7000, Thermo Fisher Scientific, MA, USA) at 19°C and 200 rpm up to 16 hours and harvested by centrifugation at 4,700 xg for 5 minutes. The cell pellets were washed by Milli-Q water once and suspended in a B-PER complete bacterial protein extraction reagent (cat # 89821, Thermo Fisher Scientific, MA, USA) to disrupt cell wall. Crude cell extract was collected by 5 minutes of centrifugation at 17,000 xg for further experiments.

#### p-nitrophenyl acetate (pNPA) assay

Deacetylation activity of the seven O-deacetylases was screened by a colorimetric pNPA assay on a 96-well plate. The reaction buffer contained 50 mM Tris-HCl at pH 6.8 and 10 mM pNPA in 200 μL of total reaction volume. The reaction started by adding crude cell extracts at 2X, 4X, and 8X dilutions and then an absorbance 405nm was monitored in a microplate reader at 50°C for 10 minutes. Specific activity (ΔAbs405/min/mg protein) was calculated based on the protein concentration determined by the Bradford assay and the measured kinetics in a linear range.

#### High-throughput screening of isobutyl acetate hydrolysis

pH change of the reaction solution was measured with a pH meter (Hach, CO, USA) and strip (cat # 13-640-502, Thermo Fisher Scientific, MA, USA) for a high-throughput screening of esterases. The reaction solution contained 20 mM Tris-HCl at pH 7.4 and 20 mM of esters in 1 mL total volume. The reaction started by adding crude cell extracts followed by 12 hours incubation at 55°C in an oven. The same volume of water was added to the negative control instead of crude cell extract. For the positive control, a single bead of Novozyme 435 (Novozymes, Denmark) was added to the reaction solution. After the reaction, the solution was centrifuged at 17,000 xg for 3 minutes and the supernatant was sampled for the pH measurement. HPLC analysis was used to verify the hydrolysis of isobutyl acetate into isobutanol and acetate.

### Bioinformatic analysis

#### Protein structure homology modeling

The homology model of Clo1313_0613 was generated using the Swiss-Model software (44) and visualized using the Molecular Operating Environment software (MOE, version 2019.01). The 3D structure was energy minimized with the Amber10: EHT force, and the binding pocket was searched using the ‘Site Finder’ tool in MOE.

#### Multiple sequence alignment and phylogenetic tree construction

Identified esterases were sequence aligned with the homolog proteins from different organisms. The representative homologous proteins of Clo1313_0613 and Clo1313_0693 were identified from the Carbohydrate Active Enzymes (CAZy) database (http://www.cazy.org/) along with BLAST search (26, 45) and their sequences were retrieved from the NCBI protein sequence repository. Their domain sequences were aligned by ClustalW (46) in Mega7 (47). Phylogenetic trees were built based on the aligned sequences using the maximum likelihood method with 1,000 bootstrap replicates. A 40% bootstrap confidence level cutoff was selected.

### Isobutyl acetate production

#### Cellulose fermentation

Tube-scale cellulose fermentation was performed as previously described (16). Briefly, 20 g/L of Avicel PH-101 was used as a sole carbon source in a 16 mL culture volume. 0.8 mL of overnight cell culture was inoculated in 15.2 mL of C-MTC medium, and 4 mL hexadecane was added in the anaerobic chamber. Each tube contained a small magnetic stirrer bar to homogenize cellulose and the culture was incubated in a water bath connected with a temperature controller set at 55°C and a magnetic stirring system. Following pH adjustment with 70 μL of 5 M KOH injection, 800 μL of cell culture and 200 μL of hexadecane layer were sampled every 12 hours.

#### Isobutanol-supplemented cellulose fermentation

The isobutanol-supplemented cellulose fermentation followed the same procedure as described above with the addition of 5 g/L isobutanol at 30 hours when the culture reached the stationary phase. The cell culture was sampled every 12 hours with intermittent pH adjustment.

### Analytical methods

#### Cell growth measurement

Cell growth in cellobiose media was measured by optical density (OD) with a spectrophotometer (Spectronic 200+, Thermo Fisher Scientific, MA, USA) at 600 nm wavelength. For the cellulose fermentation experiments, cell growth was monitored by measuring pellet protein determined by the Bradford assay (16).

#### Residual cellulose quantification

Residual cellulose quantification followed the phenol-sulfuric acid method with slight modification (16). Standard curves were determined by using Avicel PH-101 at concentrations of 20 g/L, 10 g/L, 5 g/L, 1 g/L, 0.5 g/L, and 0.1 g/L for every experiment.

#### High-performance liquid chromatography (HPLC) analysis

HPLC system (Shimadzu Inc., MD, USA) was used to quantify extracellular metabolites. 800 μL of samples were centrifuged at 17,000 xg for 3 minutes followed by filtering through 0.2 micron filters. The samples were run with 10 mN H_2_SO_4_ at 0.6 mL/min on an Aminex HPX-87H (Biorad Inc., CA, USA) column at 50°C. Refractive index detector (RID) and ultra-violet detector (UVD) at 220 nm were used to determine concentrations of sugars, organic acids, and alcohols.

#### Gas chromatography coupled with mass spectroscopy (GC/MS) analysis

GC (HP 6890, Agilent, CA, USA) equipped with a MS (HP 5973, Agilent, CA, USA) was used to quantify esters. A Zebron ZB-5 (Phenomenex, CA, USA) capillary column (30 m x 0.25 mm x 0.25 μm) was used with helium as the carrier gas at a flow rate of 0.5 mL/min. The oven temperature program was set as follows: 50°C initial temperature, 1°C/min ramp up to 58°C, 25°C/min ramp up to 235°C, 50°C/min ramp up to 300°C, and 2-minutes bake-out at 300°C. 1 μL sample was injected into the column with the splitless mode at an injector temperature of 280°C. For the MS system, selected ion mode (SIM) was used to detect and quantify esters with the parameters described previously (16). As an internal standard, 10 mg/L n-decane were added in initial hexadecane layer and detected with m/z 85, 99, and 113 from 12 to 15 minute retention time range.

## ACKNOWLEDGMENTS

This research was financially supported by the Center for Bioenergy Innovation (CBI), the U.S. Department of Energy (DOE) Bioenergy Research Centers funded by the Office of Biological and Environmental Research in the DOE Office of Science. PNN was supported by an undergraduate research internship scholarship at The University of Tennessee Knoxville. The authors would like to acknowledge the Center of Environmental Biotechnology at UTK for using the GC/MS instrument.

